# The TCA cycle and pentose phosphate pathway are linked to lipid droplet expansion in nitrogen-starved *Lipomyces starkeyi* cells

**DOI:** 10.64898/2026.05.27.728067

**Authors:** Rio Kubo-Sato, Genki Sato, Nobuyuki Okahashi, Fumio Matsuda, Koji Okamoto

**Affiliations:** Graduate School of Frontier Bioscience, The University of Osaka, 1-3 Yamadaoka, Suita, Osaka, 565-0871, Japan; Department of Bioinformatic Engineering, Graduate School of Information Science and Technology, The University of Osaka, 1-5 Yamadaoka, Suita, Osaka 565-0871, Japan; Department of Biotechnology, Osaka University Shimadzu Analytical Innovation Research Laboratory, Graduate School of Engineering, The University of Osaka, 2-1 Yamadaoka, Suita, Osaka 565-0871, Japan; Industrial Biotechnology Initiative Division, Institute for Open and Transdisciplinary Research Initiatives, The University of Osaka, 2-1 Yamadaoka, Suita, Osaka 565-0871, Japan

**Keywords:** Lipid droplet, TCA cycle, Pentose phosphate pathway, Mitochondria, Oleaginous yeast

## Abstract

In eukaryotic cells, lipid droplets (LDs) play critical roles in storing energy, preventing lipotoxicity, promoting membrane biogenesis, and regulating stress responses, contributing to the maintenance of cellular homeostasis. Similar to white adipocytes under overnutrition, the oleaginous yeast *Lipomyces starkeyi* cells form a single giant LD during nitrogen deprivation. Under the same conditions, mitochondria form elongated tubules and sheets in a close proximity to a giant LD, although the significance of this mitochondria-LD proximity remains unclear. Here, we show that inhibition of fatty acid synthesis leads to strong suppression of LD expansion in nitrogen-starved *L. starkeyi* cells. Metabolomics analysis reveals that the TCA cycle intermediates including citric acid, a key precursor for fatty acid synthesis, decrease in cells undergoing LD expansion. In contrast, the pentose phosphate pathway intermediates increase in a manner dependent on fatty acid synthesis. Inhibition of the pentose phosphate pathway, which generates NADPH, a key electron donor for fatty acid synthesis, strongly suppressed LD expansion. Surprisingly, nitrogen-starved *L. starkeyi* cells also accumulate carnitine, a critical carrier that mediates transport of fatty acids to mitochondria and accelerates beta-oxidation for energy production. Our findings raise the possibility that, under nitrogen starvation, *L. starkeyi* cells activate fatty acid synthesis with citric acid and NADPH from the TCA cycle and the pentose phosphate pathway, respectively, thereby facilitating energy production and storage concurrently via the mitochondria-LD proximity.

## Introduction

Energy storage is one of the essential anabolic events contributing to cell homeostasis, as it coordinates a balance between energy production and consumption. In almost all eukaryotes, cellular energy is stored primarily as neutral lipids that are enclosed by a phospholipid monolayer. This energy reservoir is called lipid droplet (LD), a spherical compartment whose biogenesis is established through a myriad of carbohydrate metabolic reactions and regulated with multiple interorganellar communications. LDs are highly dynamic, such that their number and size fluctuate according to nutritional status, and their expansion-contraction cycle is synchronized with cellular metabolism. In addition, they are multitasking organelles acting in storing excess carbon, maintaining lipid homeostasis, and govering stress responses (*5*, *7*). Defects in lipid droplet biogenesis are associated with a variety of disorders including metabolic syndrome, neurodegeneration, and inflammation, underscoring the physiological relevance. Although numerous studies reveal that formation of a single large lipid droplet, which is typically appreciated in white adipocytes, is beneficial for efficient fat storage and lowered lipotoxicity, how cells promote drastic lipid droplet expansion remains enigmatic.

In contrast to most of yeast species harboring several small LDs, the oleaginous yeast *Lipomyces starkeyi* can produce a single giant LD that occupies over 60% of the total cell volume, with stored lipids reaching up to 65-85% of its dry weight (*1*, *2*, *3*). In *L. starkeyi* cells, nitrogen starvation triggers this drastic LD expansion, whereas it does not induce autophagy-related processes (*11*). Strikingly, under the same conditions, mitochondria remodel their shape from fragments and rods to tubules and sheets, localizing in a close proximity to a giant LD (*11*). Currently, it remains to be investigated if there is a metabolic exchange between the two organelles, thereby establishing efficient energy storage and/or production. In addition, how nitrogen-starved *L. starkeyi* cells undergo LD expansion has yet to be elucidated.

In this study, we sought to clarify metabolic processes crucial for LD expansion in *L. starkeyi* cells under nitrogen starvation. To this end, we performed fluorescence microscopy to observe LDs and mitochondria, and mass spectrometry to analyze intracellular metabolites. We attempted to inhibit de novo synthesis of fatty acid (FA), a precursor to generate neutral lipids to form LDs, and found that FA synthesis is critical for LD expansion. Our results showed a decrease in citric acid cycle intermediates and an increase in pentose phosphate pathway intermediates. Inhibiting fatty acid synthesis suppressed lipid droplet expansion and increased TCA cycle metabolites, including citrate, a key precursor for fatty acid synthesis. Interestingly, inhibiting the pentose phosphate pathway, which is activated under nitrogen deficiency and is essential for generating NADPH (the electron donor required for fatty acid synthesis), inhibited lipid droplet expansion. These results suggest that, under nitrogen-deficient conditions, NADPH and citrate, supplied by the pentose phosphate pathway and the TCA cycle, respectively, promote lipid droplet expansion.

## Materials and Methods

### Yeast strains and growth conditions

The *Lipomyces starkeyi* strain used in this study was derived from a strain cultivated using standard genetic and molecular biological techniques (*10*). *Lipomyces starkeyi* NRRL Y-1388 (provided by the RIKEN BioResource Center) was cultured at 26°C in YPD medium (1% yeast extract, 2% peptone and 2% dextrose) or complete medium SDCA (0.17% yeast nitrogen base without amino acids and ammonium sulphate, 0.5% ammonium sulphate with 0.5% casamino acids containing 2% dextrose) supplemented with necessary amino acids and nucleotide bases. For nitrogen depletion, cells were pregrown to mid-log phase in SDCA and transferred to SD-N medium (0.17% yeast nitrogen base without amino acids and ammonium sulphate plus 2% dextrose). For carbon depletion, cells were pregrown to mid-log phase in SDCA and transferred to SN-D medium (0.17% yeast nitrogen base without amino acids and ammonium sulphate and ammonium sulphate, 0.5% ammonium sulphate with 0.5% casamino acids).

### Fluorescence microscopy

*Lipomyces starkeyi* cells were observed using a Pulse-SIM BZ-X800 microscope (Keyence) equipped with a 100 ×objective lens (CFI Apochromat TIRF 100XC Oil, NA: 1.49; Nikon), filter sets for GFP and mCherry (BZ-X filter GFP and BZ-X filter TRITC, respectively; Keyence) and an optical sectioning module (BZ-H4XF; Keyence).

### Quantification of survival rate using Phloxine B

Survival rate measurement by Phloxine B staining was performed as follows. First, 1 μL of a 1 mg/mL Phloxine B solution was added to a sample of *L. starkeyi* equivalent to 1 OD600 unit, and the mixture was left at room temperature for 5 minutes. The supernatant was then removed by centrifugation, and the pellet was resuspended and washed with Milli-Q water. After repeating this centrifugation and Milli-Q water wash twice, the pellet was resuspended in 25 μL of Milli-Q water. A 2 μL aliquot of this suspension was used as the sample for fluorescence microscopy observation. For Phloxine B staining, cells stained red throughout were determined to be dead cells. For each sample, images of five fields of view were acquired under a fluorescence microscope, and the number of live cells and dead cells within each field of view was counted. The survival rate (%) for each field of view was calculated as the number of live cells ÷ total number of cells × 100. The average survival rate from the five fields of view was then used as the survival rate for that sample.

### Lipid droplet assay

*Lipomyces starkeyi* cells pregrown in SDCA medium were transferred to SD-N media, cultured at 26℃, collected at the 0-, and 24-h time points, and observed under a fluorescence microscope. All experiments were repeated three times independently. More than 30 cells were selected randomly from microscopic images for different culture conditions at each time point. The number of cells (N) and the area of LD-specific BODIPY signals (a) was measured by ImageJ (National Institutes of Health, USA). The average area of LDs in each cell (A) was calculated by A = a/N.

### Measurement of glucose consumption

*L. starkeyi* cells were pre-cultured in SDCA medium until mid-logarithmic phase, then transferred to nitrogen-starved conditions (SD-N). Cerulenin was added at a final concentration of 45 μM to samples where lipid droplet expansion was induced. As a negative control, cells were similarly pre-cultured and then transferred to SD-N containing the same volume of DMSO as in the Cerulenin-added group. The culture supernatants were collected, and glucose levels were measured. For glucose quantification, a glucose measurement kit (Dojin Chemical Laboratory) was used. The glucose in the cell culture supernatant was detected by measuring the absorbance of WST-F, which developed color in proportion to the glucose amount.

### Metabolome analysis

*L. starkeyi* cells were recovered from the culture medium using a filter method (PTFE membrane filter: pore size 0.45 μm, diameter 47 mm; Omnipore, Merck Millipore, Kenilworth, NJ, USA). For SDCA culture, cells corresponding to OD600 ≈ 10 were recovered. For SD-N culture, cells corresponding to an OD600 ≈ 5 were recovered and divided between two filters. Cells were immersed in 1.6 mL methanol containing internal standards (20 μM D-camphor sulfonic acid, Sigma-Aldrich; 20 μM D-norvaline, TCI) and stored at −80°C. Intracellular metabolites were extracted using the methanol/chloroform/water extraction method by adding 640 μL of Milli-Q water and 1.6 mL of chloroform. The resulting mixture was centrifuged (3,700 × g, 4°C, 20 min), and 250 μL of the supernatant was aliquoted into 6 or 3 × 1.5 mL tubes and dried under vacuum using a SpeedVac (Thermo Fisher Scientific). For SD-N medium, two filter Metabolite data were acquired by ion-pair liquid chromatography-triple quadrupole mass spectrometry (LC-MS/MS) and gas chromatography-quadrupole mass spectrometry (GC-MS), performed as previously reported (*14*). For GC-MS analysis, 100 μL of hydrochloric acid methoxamine in pyridine solution (40 mg/mL) and 100 μL of N-methyl-N-(tert-butyldimethylsilyl) trifluoroacetamide (MTBSTFA) containing 1% tert-butyl dimethyl chlorosilane (TBDMCS) to derivatize the dried sample. All data were processed using LabSolution software (Version 5.1, Shimadzu Corporation).

### Statistical Analysis

Results are presented as mean ± standard deviation (SD). Statistical testing was performed using Welch’s t-test. For metabolome analysis, multiple testing was corrected using the Benjamini-Hochberg method. The significance level was set at p<0.05. Calculations were performed using Microsoft Excel (Microsoft Office 2020) and JMP Student Edition 19 (JMP Statistical Discovery LLC).

## Results

### Lipid droplets begin to enlarge around 12 hours under nitrogen starvation

To determine the timing of lipid droplet expansion initiation in *L. starkeyi*, cells expressing the mitochondrial substrate-targeted probe mito-DHFR-mCherry, a mitochondrial substrate-targeted probe consisting of dihydrofolate reductase (DHFR) and mCherry, were cultured in complete nutrient medium (SDCA) until OD600≈1. Samples were then transferred to nitrogen-depleted medium (SD-N: –N), which induces LD expansion, and the progression of lipid droplet expansion was observed over time. As a negative control, samples cultured for the same duration in complete nutrient medium (SDCA), a condition where lipid droplets do not expand, were compared and observed. Fluorescence microscopy was used to quantify LD size using BODIPY 493/503 staining and to observe mitochondrial morphology via mito-DHFR-mCherry (Figure 1A and B). At the start of culture, LD size was 1.0 ± 0.2 μm² in SD-N and 0.9 ± 0.0 μm² in SDCA. No significant difference was observed between the two at the start of culture (*p* = 0.18). After 6 hours of culture, SD-N LD size increased to 1.3 ± 0.2 μm², while SDCA LD size was 1 ± 0.1 μm². A significant difference was observed between the SD-N sample at 6 hours post-incubation and the SDCA sample (*p* = 0.02). After 12 hours of incubation, SD-N reached 2.34 ± 0.3 μm², while SDCA reached 1 ± 0.2 μm². A significant difference was observed between the SD-N sample at 12 hours post-incubation and the SDCA sample (*p* = 0.00). At 18 hours post-incubation, SD-N measured 6.15 ± 0.9 μm², while SDCA measured 0.9 ± 0.05 μm². A significant difference was observed between the SD-N sample at 18 hours post-incubation and the SDCA sample (*p* = 0.00). At 24 hours post-incubation, SD-N was 9.43 ± 1.4 μm², and SDCA was 1.0 ± 0.19 μm². A significant difference was observed between the SD-N sample at 24 hours post-incubation and the SDCA sample (*p* = 0.00). This indicates that lipid droplet expansion begins 12 hours after the start of incubation under nitrogen-starved conditions (Figure 1A and B, C).

**Fig. 1.**
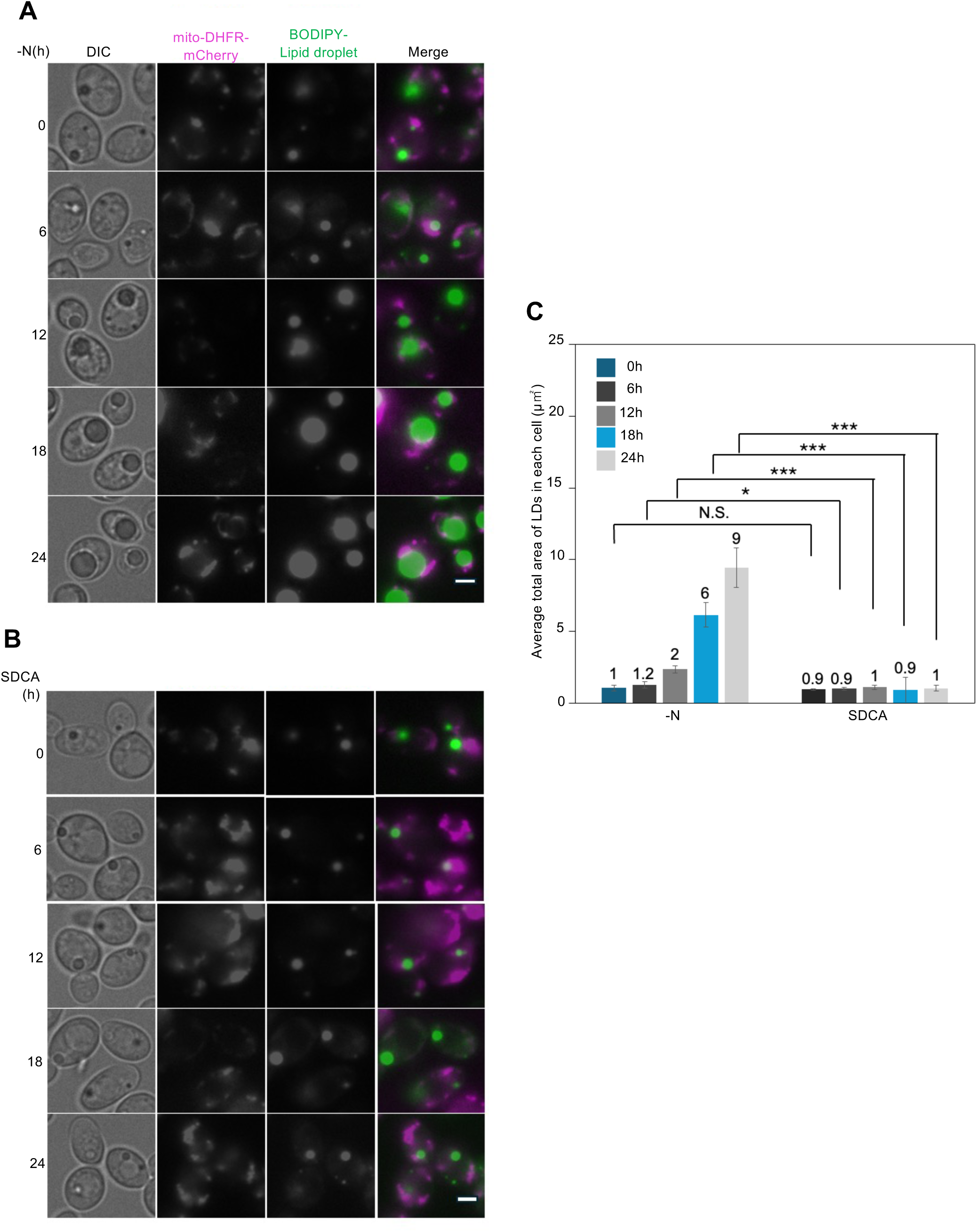
*L. starkeyi* cells under nitrogen starvation conditions exhibit lipid droplet expansion 12 h after the start of culture. (A and B) Cells expressing mito-DHFR-mCherry were grown under nitrogen (–N) or Nutrient-rich (SDCA) starvation. Mitochondrial shapes were analysed at the indicated time points using a fluorescence microscope. (C) The average total area of LDs in each cell was quantified by the area of BODIPY signals. Scale bar, 5 μm.

**Fig. 2.**
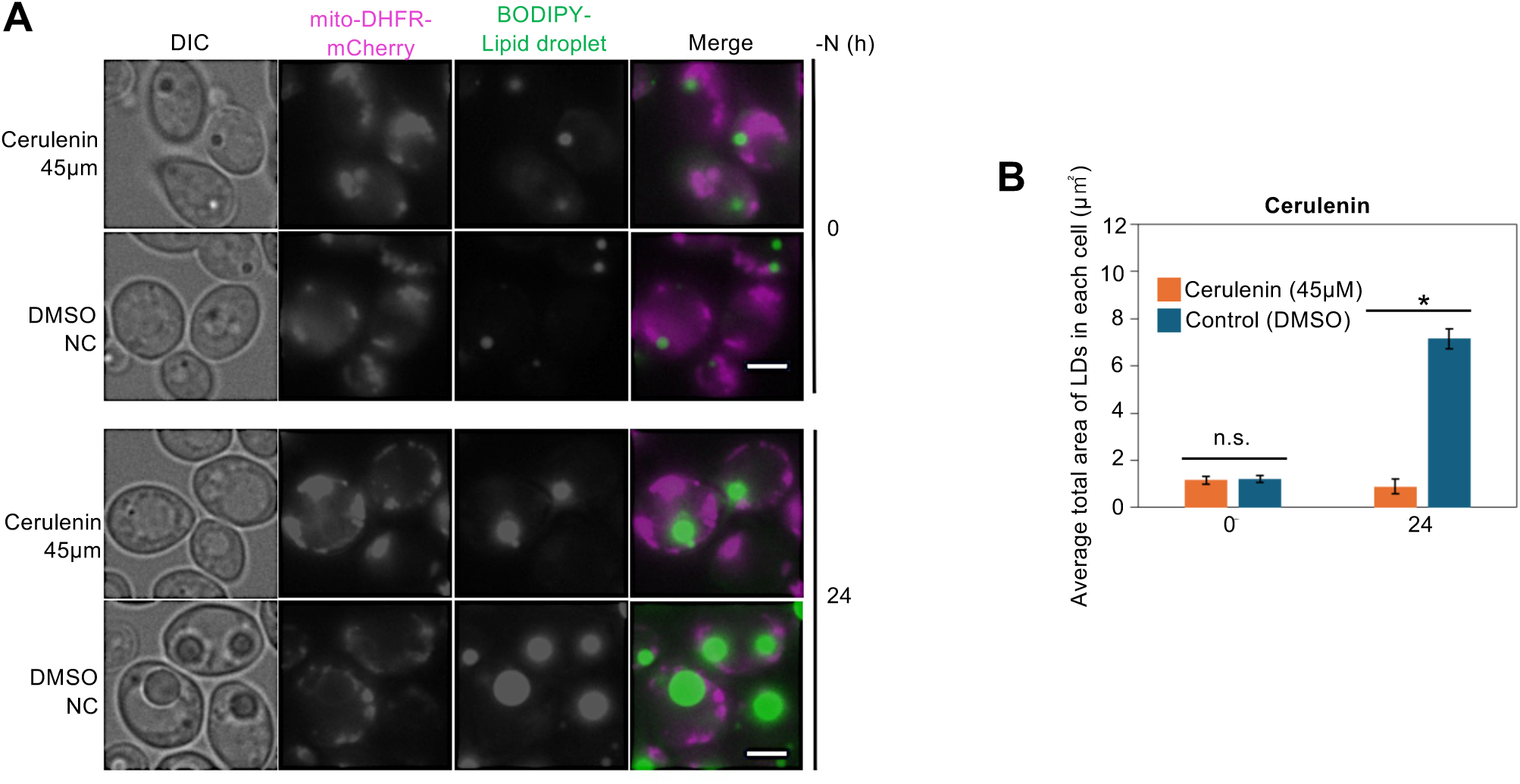
Adding cerulenin to *L. starkeyi* cells under nitrogen-starvation conditions suppresses lipid droplet expansion. (A) Cells expressing mito-DHFR-mCherry were pre-cultured in SDCA and then transferred to nitrogen-starved medium (SD–N) with or without cerulenin (45 μM). Mitochondrial morphology was analysed at 0 and 24 h using fluorescence microscopy. (B) Lipid droplets (LDs) were visualized by BODIPY staining. The average total LD area per cell was quantified from fluorescence images using ImageJ (≥30 cells per condition). Statistical significance was assessed by Welch’s t-test. *: *p*<0.05, N.S.: *p*>0.05.

### Cerulenin administration does not significantly affect glucose consumption

To determine whether cerulenin-induced inhibition of fatty acid synthesis and suppression of lipid droplet expansion in *L. starkeyi* cells affects intracellular energy production (glucose consumption), glucose was quantified in the culture supernatant of samples treated with cerulenin. Cells visualized with mito-DHFR-mCherry for mitochondria and BODIPY for lipid droplets were pre-cultured with SDCA until mid-log phase. They were then transferred to the nitrogen-depleted medium SD-N and cerulenin was added simultaneously. Additionally, as a negative control, an equal volume of DMSO was added instead of the inhibitor. For glucose quantification, a glucose measurement kit (Dojin Chemical Laboratory) was used. Glucose in the cell culture supernatant was detected by measuring the absorbance of WST-F (formazan) produced in proportion to the glucose amount. Analysis of glucose concentration in culture supernatant under nitrogen starvation revealed that the 0-hour sample contained 127.0 ± 7.8 mM, while the 24-hour sample contained 104.2 ± 8.5 mM, showing a significant decrease between 0 and 24 hours (p=0.012). Meanwhile, the glucose concentrations in the 24-hour sample and the negative control were 104.2 ± 8.5 mM and 110.8 ± 3.3 mM, respectively, showing no significant difference between the two (p=0.0153) (Supplementary Figure 1A and B).

We also examined the effect of cerulenin addition on cell growth. The specific growth rate was calculated from the OD_600_ after 24 hours of nitrogen starvation and the initial OD. Error bars indicate standard deviation. Significant differences between culture conditions for SDCA were calculated using Dunnet’s t-test. The specific growth rate of SDCA cells was 0.14 ± 0.00 h⁻¹. The specific growth rate of SDCA cells supplemented with cerulenin was 0.09 ± 0.00 h⁻¹. The specific growth rate of SD-N cells was 0.09 ± 0.00 h⁻¹. The specific growth rate of cells in SD-N supplemented with cerulenin was 0.08 ± 0.00 h⁻¹. A significant difference was observed between cells in SDCA and those in SDCA supplemented with cerulenin (p = 0.00). Similarly, a significant difference was observed between cells in SDCA and those in SD-N (p = 0.00). However, no significant difference was observed between SD-N and SD-N cells supplemented with cerulenin (p = 0.00) (Supplementary Figure 1C). These results suggest that cerulenin supplementation under nitrogen starvation suppresses lipid droplet expansion but does not significantly affect cellular energy metabolism or growth.

### Metabolomic changes in TCA cycle and pentose phosphate pathway intermediates were characterized lipid droplet formation

We subjected *L. starkeyi* cells to metabolomics analysis to elucidate the progression of lipid droplet expansion and fluctuations in intracellular metabolism under conditions in which fatty acid synthesis was inhibited. The cells, which were visualized with mito-DHFR-mCherry for the mitochondria and BODIPY for the lipid droplets, were pre-cultured in SDCA medium until the mid-log phase. Then, they were transferred to nitrogen-deficient SD-N medium that was supplemented with cerulenin. Three types of samples were used for metabolite analysis: cells cultured in cerulenin-supplemented and supplemented SD-N medium and cells cultured in SDCA medium with a nitrogen source supplemented with essential amino acids and nucleic acids. The cells were recovered using a filtration method. The metabolites were extracted with methanol-chloroform, and two types of targeted LC-MS/MS and GC-MS analyses were performed. As a result, a total of 173 intracellular metabolites were quantified.

A principal component analysis (PCA) of metabolite levels revealed differences in culture conditions. The first principal component (PC1) distinguished the presence or absence of a nitrogen source, and the second principal component (PC2) distinguished the addition or non-addition of cerulenin (Fig. 3A). In other words, the metabolite level data reflected the characteristics of the cellular state under each culture condition. Examining the contribution of metabolite levels to each principal component (Fig. 3B) revealed that metabolites with large positive contributions to PC1 were amino acids and their biosynthetic intermediates, as well as TCA cycle intermediates. Conversely, metabolites with large negative contributions were intermediates downstream of glycolysis and metabolites from the non-oxidative pentose phosphate pathway. These results are consistent with previous reports showing decreased intracellular amino acid levels during lipid yeast cultivation under nitrogen deficiency (*15*). Metabolites with large positive contributions to PC2 were phospholipid-related metabolites and TCA cycle intermediates. Conversely, metabolites with large negative contributions included glycolytic and trehalose biosynthesis intermediates and nucleic acid-related metabolites supplied by the non-oxidative pentose phosphate pathway. Specifically, SD-N culture, in which lipid droplet expansion occurs, was negatively separated from other culture conditions on both the first and second principal components. Changes in metabolite levels during lipid droplet expansion included the accumulation of downstream glycolytic and non-oxidative pentose phosphate pathway intermediates, as well as a decrease in TCA cycle intermediates.

**Fig. 3.**
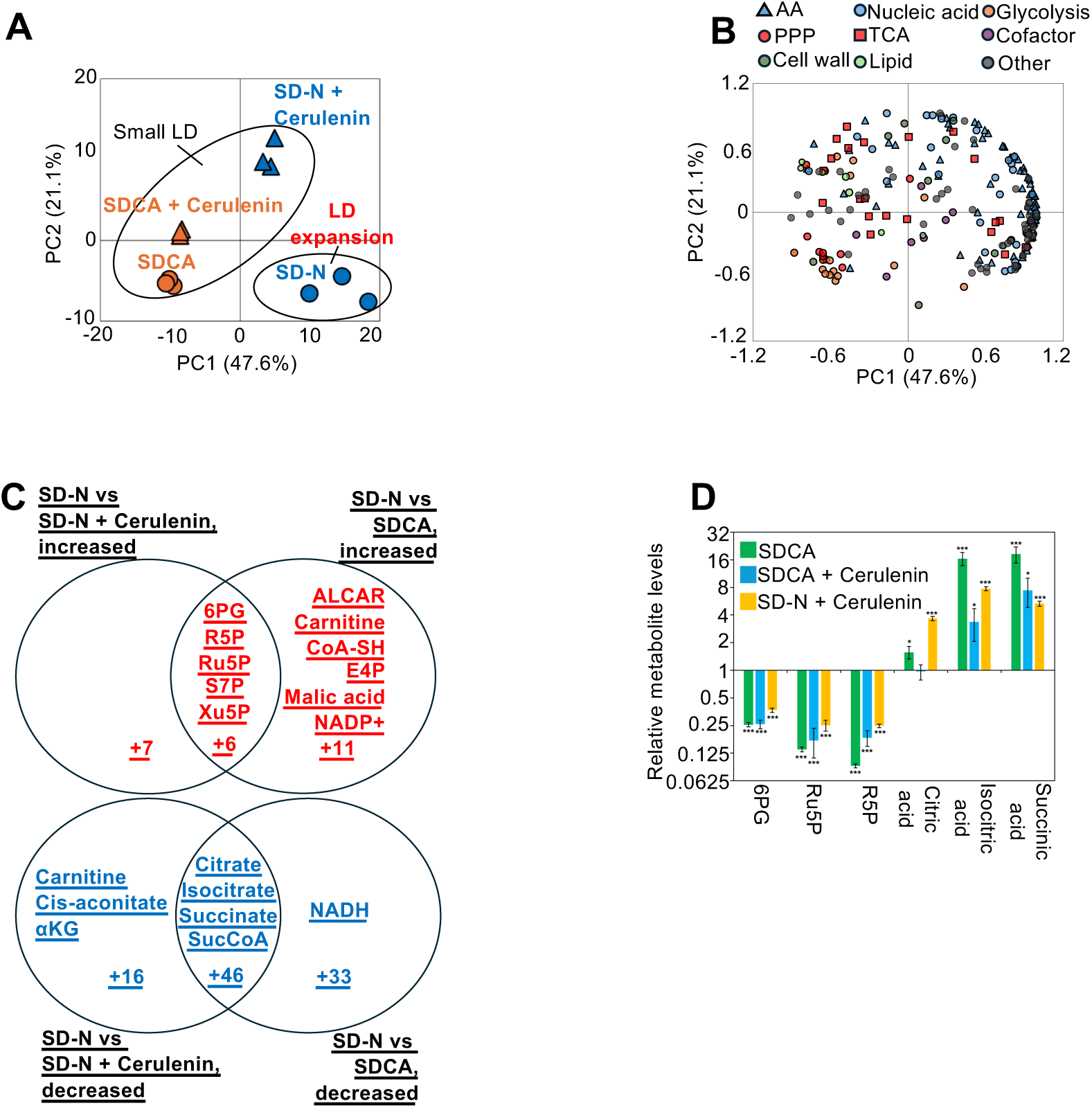
Under nitrogen-starvation conditions, the levels of metabolites in the pentose phosphate pathway increased, while the levels of metabolites in the citric acid cycle decreased. (A) Cells expressing mito-DHFR-mCherry were pre-cultured in SDCA medium, then transferred to nitrogen-depleted medium (SD–N) with or without cerulenin (45μM) and complete nutrient medium (SDCA). (B) Intracellular metabolites were extracted and analysed by targeted LC–MS/MS and GC–MS. In total, 173 metabolite levels were used for principal component analysis (PCA). PCA score plot for culture condition was shown. (B) Loading plot of major two principal components. The color of plots shows the functional annotation of the metabolites. (C) Comparison of significantly altered metabolites between culture conditions. 6PG: 6-phosphogluconate, R5P: ribose 5-phosphate, Ru5P: ribulose 5-phosphate, S7P: Sedoheptulose 7-phosphate, Xu5P: Xylulose 5-phosphate, ALCAR: Acetyl-L-carnitine, CoA-SH: Coenzyme A, E4P: Erythrose 4-phosphate, αKG: alpha-ketoglutarate, SucCoA: succinyl-CoA. (D) Relative metabolite levels are shown in relation to lipid droplet expansion. Statistical significance is indicated as \**p* < 0.05, \*\**p* < 0.01, \*\*\**p* < 0.001.

To analyze the metabolites that showed significant variation between the culture conditions, a volcano plot was created, and a Venn diagram was generated from the obtained data. To eliminate the effects of stress conditions (nitrogen starvation and cerulenin addition), the SD-N + cerulenin condition was compared with the SDCA condition, and metabolites showing variation were excluded. As a result, seven metabolites were significantly increased under the SD-N condition, where lipid droplet expansion occurs, compared to the SDCA condition, where lipid droplet expansion is not induced. Under the SD-N + cerulenin condition, 17 metabolites showed significant increases. Significant increases were detected in 11 metabolites common to both conditions. Under the SD-N condition, where lipid droplet hypertrophy occurs, a significant decrease in 19 metabolites was detected compared to the SDCA condition, where lipid droplet hypertrophy is not induced. Under the SD-N + cerulenin condition, a significant decrease in 34 metabolites was detected. A significant decrease in 50 metabolites common to both conditions was detected (Figure 3C). No particularly noteworthy metabolites were identified among the increased metabolites under the SD-N + cerulenin depletion condition when considering cerulenin stress. Under the SDCA condition, carnitine and its related metabolite, ALCAR, increased. These metabolites are associated with carnitine metabolism, which transports fatty acids into mitochondria and converts them to ATP. Other increased metabolites included CoA-SH, which is crucial for acyl group activation and transfer during energy production, lipid/amino acid synthesis, and degradation; E4P, an intermediate in the non-oxidative pentose phosphate pathway; malate, an intermediate in the citric acid cycle; and NADP+, the level of which fluctuates with increased NADPH consumption. The intermediates of the non-oxidative pentose phosphate pathway that increased under both conditions were 6PG, R5P, Ru5P, S7P, and Xu5P. Under SD-N+ cerulenin conditions, levels of carnitine and the citric acid cycle intermediates cis-aconite and α-ketoglutarate were significantly reduced. This aligns with the requirement of phospholipids as substrates and NADPH as a reducing agent, which are supplied by the pentose phosphate pathway, for lipid synthesis in Saccharomyces cerevisiae (*16*). Conversely, under SDCA conditions, NADH levels were significantly reduced. This suggests rapid NADH oxidation under complete medium conditions, indicating active energy production. Metabolites that were significantly reduced under both conditions included citrate, isocitrate, succinate, and succinate CoA, which are all intermediates of the citric acid cycle (Fig. 3C). The characteristic metabolic changes that occur during lipid droplet expansion are considered to be the accumulation of 6PG, Ru5P, and R5P, as well as the depletion of intermediates such as citrate and α-ketoglutarate on the decarboxylation side of the citric acid cycle (Fig. 3D).

Furthermore, the molar ratios of coenzymes were analyzed under each culture condition (Fig. 4A). The ATP/ADP and NADH/NAD ratios decreased in the following order: SDCA, SD-N, and SD-N + cerulenin. Similar changes have been reported in yeast cells cultured in the presence of electron transport chain uncouplers or complex III inhibitors. This suggests a state of reduced efficiency in the mitochondrial electron transport chain(*17*). Additionally, no significant differences were observed in oxidized glutathione levels or NADPH/NADP ratios when comparing SD-N and SD-N + cerulenin cultures. This indicates that intracellular oxidative stress did not increase. Therefore, under nitrogen starvation conditions, inhibiting fatty acid synthesis may reduce mitochondrial respiratory efficiency.

**Fig. 4.**
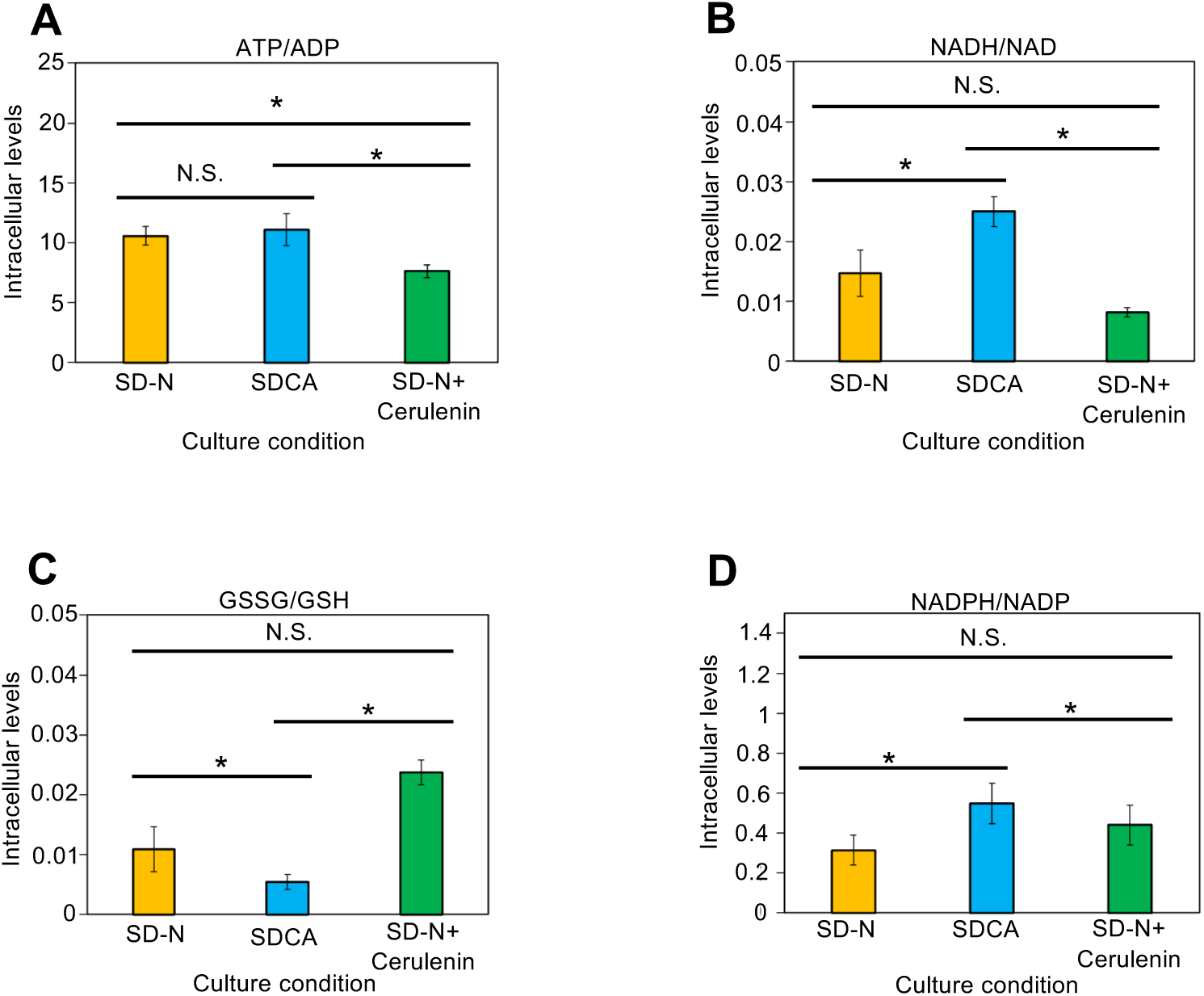
Adding cerulenin to *L. starkeyi* cells under nitrogen-limited conditions reduces intracellular ATP levels. (A) Ratio of ATP to ADP abundance under each culture condition. (B) Ratio of peak area values standardized by internal standard for NADH to NAD abundance. (C) Oxidized glutathione abundance under each culture condition. Peak area values standardized by internal standard are shown. (D) Ratio of NADPH to NADP. Shows the ratio of peak area values normalized by internal standard. Significant differences between culture conditions were calculated using Welch’s t-test, and p-values were controlled for false positives using the Holm method. *: *p*<0.05, N.S.: *p*>0.05.

### Inhibition of the pentose phosphate pathway activity blocks lipid droplet expansion

We investigated whether 6-aminonicotinamide, which causes a decrease in NADPH levels via inhibition of G6PD in the pentose phosphate pathway, could suppress the expansion of lipid droplets in *L. starkeyi* cells. For observation, cells visualized for mitochondria using mito-DHFR-mCherry and for lipid droplets using BODIPY were pre-cultured with SDCA until mid-log phase. They were then transferred to nitrogen-starved SD-N medium with simultaneous addition of 6-aminonicotinamide. As a negative control, an equivalent volume of H₂O was added. Observations were performed at the start of culture and at the 24-hour time point. For each culture condition and time point, over 30 cells were randomly selected from microscopic images. The number (n) and diameter (d) of lipid droplets, as well as the total number of cells (Nc), were counted and measured using ImageJ (National Institutes of Health, USA). The average diameter of each lipid droplet was calculated as D = ‘d. The average area of lipid droplets within each cell in the acquired fluorescence microscope images was calculated as A = π * (D/4) ² * n/Nc μm², while the DMSO-treated sample was μm². At the start of culture, no significant difference was observed between the two (*p* = 0.313). Meanwhile, after 24 hours of nitrogen starvation, the cerulenin-treated sample measured 0.9 ± 0.3 μm², while the DMSO-treated sample measured 7.1 ± 0.418 μm². A significant difference was observed between the cerulenin-added sample and the DMSO-added sample after 24 hours of nitrogen starvation (*p* = 0.00).

The sample treated with 6-aminonicotinamide showed an initial area of 1.14 ± 0.2 μm² at the start of culture, while the DMSO-treated sample showed 1.7 ± 0.3 μm². No significant difference was observed between the two at the start of culture (*p* = 0.3). However, after 24 hours of culture, the 6-aminonicotinamide-added sample measured 1.2 ± 0.2 μm², while the DMSO-added sample measured 14.1 ± 0.9 μm². A significant difference was observed between the 6-aminonicotinamide-added sample and the DMSO-added sample after 24 hours of nitrogen starvation (*p* = 0.00). That is, 6-aminonicotinamide suppressed the expansion of lipid droplets (Figure 4A). 6-Aminonicotinamide causes a decrease in NADPH levels via inhibition of G6PD in the pentose phosphate pathway. Since NADPH production increases due to activation of the pentose phosphate pathway under nitrogen-starved conditions, this suggests that the supply of reducing power derived from NADPH, which is essential for fatty acid synthesis, may be important for lipid droplet expansion.

## Discussion

To test, from a metabolic perspective, the significance of the previously reported co-localization of mitochondria and lipid droplets (LDs) in *Lipomyces starkey*i under nitrogen deprivation, we performed quantitative intracellular metabolomics under three culture conditions: a standard medium (SDCA), a nitrogen-depleted medium (SD-N), and a nitrogen-depleted medium supplemented with cerulenin (SD-N + Cerulenin), which inhibits fatty acid synthesis and suppresses LD expansion under nitrogen starvation. In total, 173 intracellular metabolites were quantified. Principal component analysis (PCA) clearly separated the culture conditions, with PC1 reflecting the presence/absence of a nitrogen source and PC2 reflecting the presence/absence of cerulenin, indicating that the obtained metabolic profiles robustly capture the cellular state under each condition (Fig. 3A).

Previous studies in *L. starkeyi* have primarily described the phenomenon that mitochondria relocate near expanding LDs under nitrogen starvation; however, the functional and causal significance of this proximity—namely, what metabolic advantage it provides—has remained unresolved. Building on this gap, our strategy was to move beyond morphology and interrogate the LD expansion state through metabolome-wide readouts, while also introducing a targeted perturbation (cerulenin) that specifically blocks fatty acid synthesis and LD expansion. This design allows us to distinguish (i) metabolic changes that represent a general nitrogen-starvation response from (ii) changes that are tightly associated with the LD expansion program itself.

As a first validation step, under nitrogen starvation (SD-N), amino acids and their biosynthetic intermediates decreased, while intermediates of the (non-oxidative) pentose phosphate pathway accumulated (Fig. 3B). This pattern is consistent with known nutritional responses in lipid-producing yeasts, supporting the validity of our metabolome analysis.

Next, to extract metabolic changes that are specific to LD expansion (rather than general stress effects associated with nitrogen starvation or cerulenin addition), we compared conditions in a way that excludes these broad effects. Under LD expansion-promoting conditions, significant increases were detected in PPP-related metabolites including 6PG and R5P, as well as Ru5P, S7P, and Xu5P. In contrast, TCA cycle intermediates—citrate, isocitrate, succinate, and succinyl-CoA—showed significant decreases (Fig. 3C and D). Because intracellular metabolite levels are shaped by the balance of production and consumption, the observed decrease in TCA intermediates could reflect not only reduced TCA cycling but also enhanced efflux or increased local consumption. Importantly, the reduction of citrate during LD expansion is consistent with a model in which mitochondria-derived citrate is exported to the cytoplasm and consumed as a substrate for fatty acid synthesis, thereby promoting LD expansion.

We further found that inhibiting fatty acid synthesis under nitrogen starvation reduced the ATP/ADP ratio and NADH/NAD ratio (Fig. 4A and D). Similar decreases have been reported in yeast cultured with uncoupling agents or respiratory chain inhibitors(*17*), suggesting that mitochondrial energy metabolism efficiency may be reduced when fatty acid synthesis is blocked under nitrogen starvation. In contrast, oxidized glutathione levels and the NADPH/NADP ratio did not significantly differ between SD-N and SD-N + Cerulenin, suggesting that elevated oxidative stress is not the primary driver of the reduced mitochondrial function. Together, these results imply that fatty acid synthesis under nitrogen starvation contributes to maintaining cellular energy status, potentially by preserving mitochondrial respiratory efficiency.

Moreover, prior studies have proposed a pathway in which NADPH can be converted into energy-producing coenzymes such as FADH2 and NADH, occurring concomitantly with cytosolic fatty acid synthesis and mitochondrial fatty acid breakdown via β-oxidation (*18*). In line with this, we observed carnitine accumulation—an indicator of fatty acid transport into mitochondria—under conditions promoting LD expansion (Fig. 5A). This finding suggests that fatty acid degradation contributes to energy production during LD expansion, raising the possibility that LD–mitochondria proximity supports not only carbon storage but also energy generation. As a future direction, we plan to inhibit β-oxidation with acrylic acid and examine its effects on LD expansion and growth under nitrogen-depleted conditions, thereby testing this proposed dual role.

**Fig. 5.**
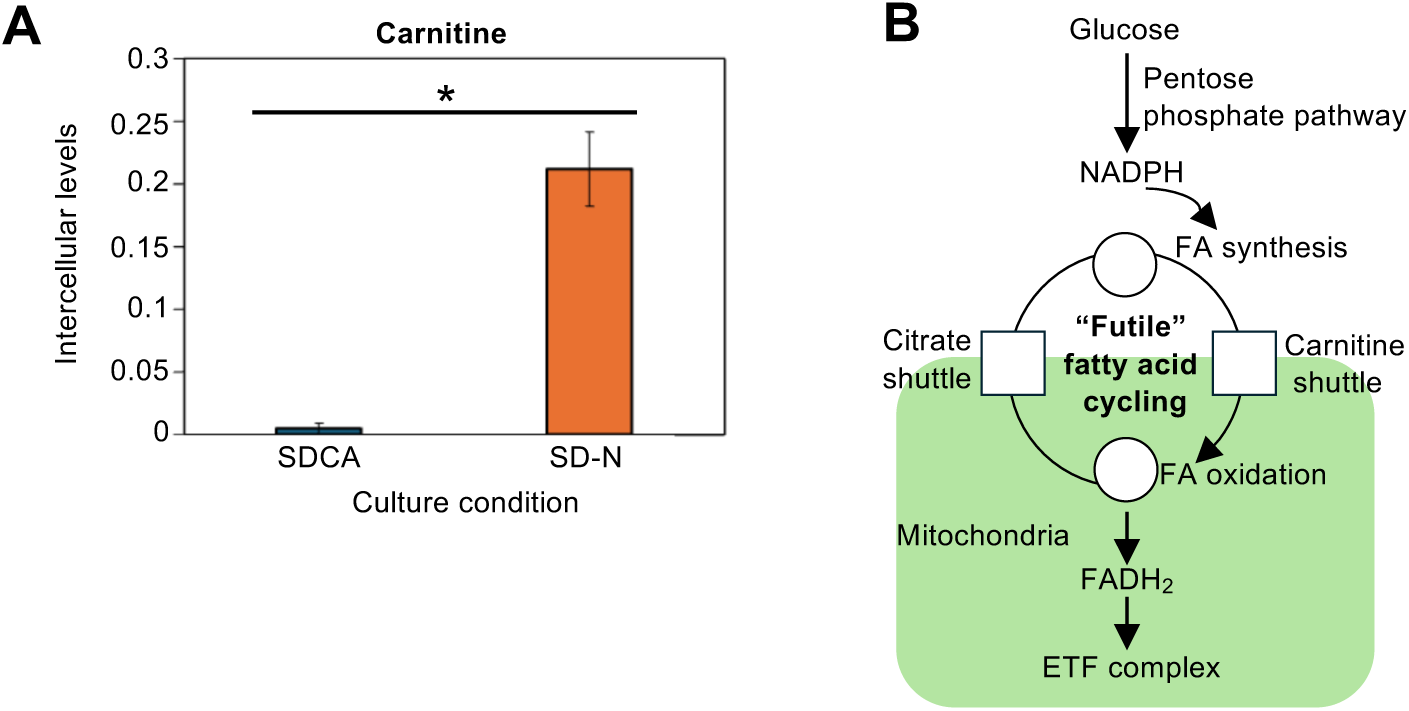
Carnitine accumulation occurs in *L. starkeyi* cells under nitrogen-limited conditions. (A) The ratio of peak area values normalized by internal standard is shown. Significant differences between culture conditions were calculated using Welch’s t-test, and p-values were controlled for false positives using Holm’s method. *: *p*<0.05, N.S.: *p*>0.05. (B) NAD+-independent glucose oxidation. Inhibition of Complex I blocks the default pathway of glucose oxidation via glycolysis and the citric acid cycle. As a result, intracellular glucose-6-phosphate is diverted to the pentose phosphate pathway. The pathway’s NADPH output is converted into FADH2 by fatty acid cycling. The resulting FADH₂ can then fuel the respiratory chain via the electron-transferring flavoprotein (ETF) complex.

**Fig. 6.**
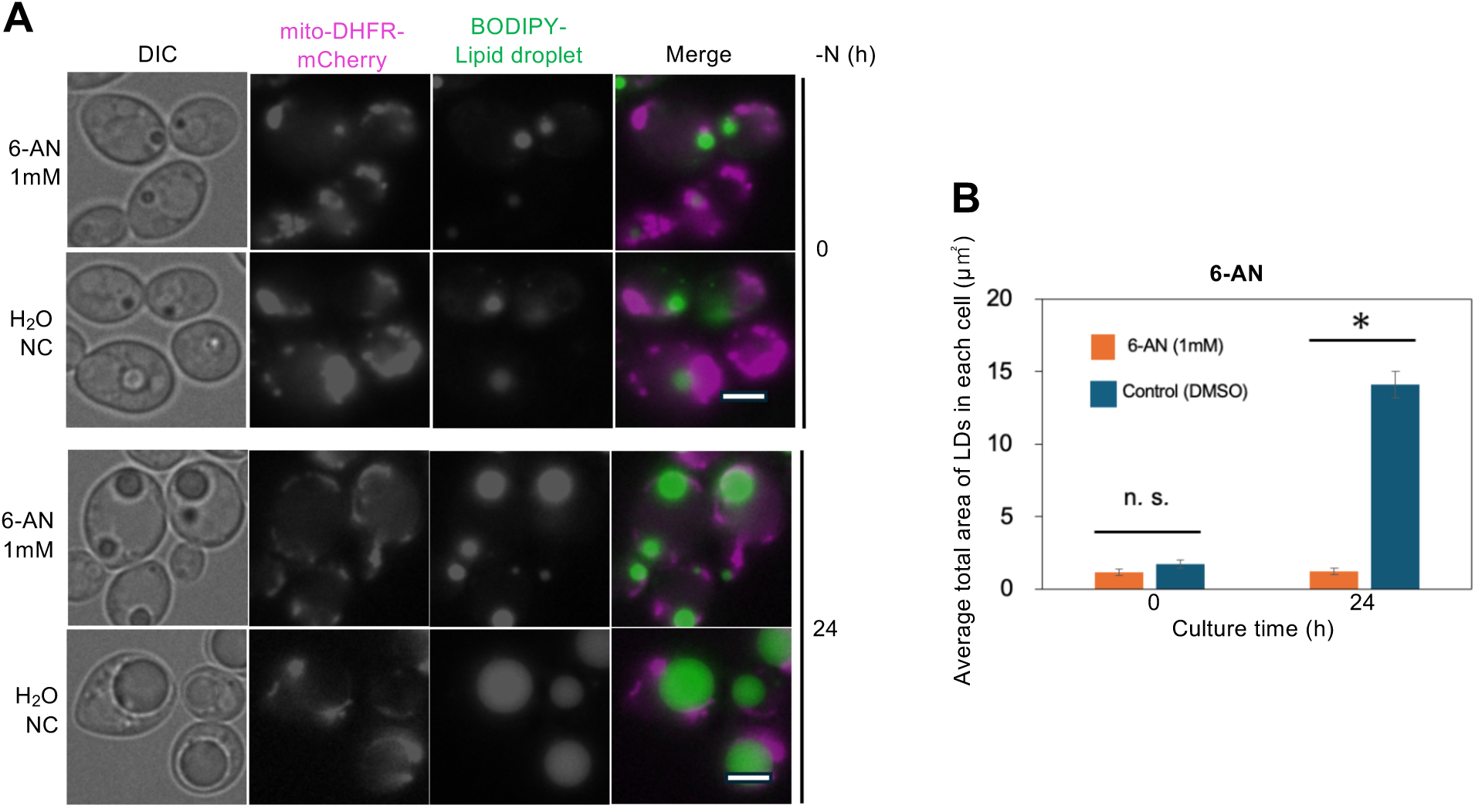
Adding 6-AN to *L. starkeyi* cells under nitrogen-starvation conditions suppresses lipid droplet expansion. (A) Cells expressing mito-DHFR-mCherry were pre-cultured in SDCA and then transferred to nitrogen-starved medium (SD–N) with or without 6-AN (1mM). Mitochondrial morphology was analysed at 0 and 24 h using fluorescence microscopy. (B) Lipid droplets (LDs) were visualized by BODIPY staining. The average total LD area per cell was quantified from fluorescence images using ImageJ (≥30 cells per condition). Statistical significance was assessed by Welch’s t-test. *: *p*<0.05, N.S.: *p*>0.05.

Taken together, our data indicate that the LD–mitochondria proximity observed in *L. starkeyi* under nitrogen starvation is not merely a passive juxtaposition for simple metabolite exchange. Rather, it likely represents an organizational arrangement that supports cellular health and intracellular energy homeostasis by coordinating (i) the pool of TCA intermediates (including cytosolic citrate availability), (ii) fatty acid synthesis that consumes these intermediates, and (iii) mitochondrial energy metabolism linked to fatty acid turnover. Because excessive accumulation of TCA intermediates in mitochondria or the cytoplasm may inhibit cell growth (*19*), channeling citrate into fatty acid synthesis and LD expansion may function as a protective mechanism to avoid metabolic imbalance. In this sense, LD–mitochondria contact may act as a specialized metabolic platform that integrates the “energy storage” and “energy production” axes during nitrogen starvation.

## Acknowledgments

This work was supported in part by Noda Institute for Scientific Research Grant (to K.O.), and the University of Osaka Transdisciplinary Program for Biomedical Entrepreneurship and Innovation (to R.K.-S.).

## References

1. McNeil BA, Stuart DT. Optimization of C16 and C18 fatty alcohol production by an engineered strain of *Lipomyces starkeyi*. J Ind Microbiol Biotechnol 45: 1–14, 2018.

2. Angerbauer C, Siebenhofer M, Mittelbach M, Guebitz GM. Conversion of sewage sludge into lipids by *Lipomyces starkeyi* for biodiesel production. Bioresour Technol 99: 3051–3056, 2008.

3. Juanssilfero AB, Kahar P, Amza RL, Miyamoto N, Otsuka H, Matsumoto H, Kihira C, Thontowi A, Yopi, Ogino C, Prasetya B, Kondo A. Effect of inoculum size on single-cell oil production from glucose and xylose using oleaginous yeast *Lipomyces starkeyi*. J Biosci Bioeng 125: 695–702, 2018.

4. Spinelli JB, Haigis MC. The multifaceted contributions of mitochondria to cellular metabolism. Nat Cell Biol 20: 745–754, 2018.

5. Olzmann JA, Carvalho P. Dynamics and functions of lipid droplets. Nat Rev Mol Cell Biol 20: 137–155, 2019.

6. Martínez-Reyes I, Chandel NS. Mitochondrial TCA cycle metabolites control physiology and disease. Nat Commun 11: 102, 2020.

7. Walther TC, Farese RV Jr. Lipid droplets and cellular lipid metabolism. Annu Rev Biochem 81: 687–714, 2012.

8. Ellis RJ. Macromolecular crowding: an important but neglected aspect of the intracellular environment. Curr Opin Struct Biol 11: 114–119, 2001.

9. Luby-Phelps K. The physical chemistry of cytoplasm and its influence on cell function: an update. Mol Biol Cell 24: 2593–2596, 2013.

10. Duan L, Okamoto K. Mitochondrial dynamics and degradation in the oleaginous yeast *Lipomyces starkeyi*. Genes Cells 26: 627–635, 2021.

11. Duan L, Togou A, Ohta K, Okamoto K. Mitochondria-giant lipid droplet proximity and autophagy suppression in nitrogen-depleted oleaginous yeast *Lipomyces starkeyi* cells. J Biochem 177: 15–25, 2025.

12. Prinz WA, Toulmay A, Balla T. The functional universe of membrane contact sites. Nat Rev Mol Cell Biol 21: 7–24, 2020.

13. Rambold AS, Cohen S, Lippincott-Schwartz J. Fatty acid trafficking in starved cells: regulation by lipid droplet lipolysis, autophagy, and mitochondrial fusion dynamics. Dev Cell 32: 678–692, 2015.

14. Nishiguchi H, Liao J, Shimizu H, Matsuda F. Novel allosteric inhibition of phosphoribulokinase identified by ensemble kinetic modeling of Synechocystis sp. PCC 6803 metabolism. Metab Eng Commun 11: e00153, 2020.

15. Pomraning KR, Kim Y-M, Nicora CD, Chu RK, Bredeweg EL, Purvine SO, Hu D, Metz TO, Baker SE. Multi-omics analysis reveals regulators of the response to nitrogen limitation in *Yarrowia lipolytica*. BMC Genomics 17: 138, 2016.

16. Zhou W, Wang Y, Zhang J, Zhao M, Tang M, Zhou W, Gong Z. A metabolic model of *Lipomyces starkeyi* for predicting lipogenesis potential from diverse low-cost substrates. Biotechnol Biofuels 14: 148, 2021.

17. Sawai A, Taniguchi T, Noguchi K, Seike T, Okahashi N, Takaine M, Matsuda F. ATP supply from cytosol to mitochondria is an additional role of aerobic glycolysis to prevent programmed cell death by maintenance of mitochondrial membrane potential. Metabolites 15: 461, 2025.

18. Abrosimov R, Baeken MW, Hauf S, Wittig I, Hajieva P, Perrone CE, Moosmann B. Mitochondrial complex I inhibition triggers NAD+-independent glucose oxidation via successive NADPH formation, “futile” fatty acid cycling, and FADH2 oxidation. GeroScience 46: 3635–3658, 2024.

19. Nishio K, Kawarasaki T, Sugiura Y, Matsumoto S, Konoshima A, Takano Y, Hayashi M, Okumura F, Kamura T, Mizushima T, Nakatsukasa K. Defective import of mitochondrial metabolic enzyme elicits ectopic metabolic stress. Sci Adv 9: eadf1956, 2023.

