## Supplemental Figures for "The TCA cycle and pentose phosphate pathway are linked to lipid droplet expansion in nitrogen-starved *Lipomyces starkeyi* cells"

**Fig. S1. The presence or absence of selenium does not result in significant differences in glucose consumption or growth.**

(A and B) The glucose levels in the culture supernatants of the cerulenin-added group and the SD-N group were calculated by measuring the absorbance of WST-FORMAZAN, which was color-developed according to the glucose amount using a quantitative kit. Significant differences in SDCA levels among the culture conditions were calculated using Dunnet's t-test. *: $p<0.05$, N.S.: $p>0.05$. (C) The specific growth rate was calculated from the OD600 value after 24-hour culture in SD-N and the initial OD600 value. Error bars indicate standard deviation. Significant differences between culture conditions relative to SDCA were calculated using Dunnet's t-test. *: $p<0.05$. N.S.: $p>0.05$.

**Fig. S2. Volcano plot showing changes in metabolite levels across culture conditions in *L. starkeyi* cells under nitrogen starvation.**

(A) Comparison between lipid droplet hypertrophy and non-hypertrophy conditions. (B) Comparison with and without cerulenin addition under nitrogen starvation. The vertical axis shows the significance of change expressed as -log10 *p*-values, while the horizontal axis shows the change in mean values expressed as log2. The vertical axis line indicates the significance level of $p = 0.05$, and the horizontal axis lines indicate changes of 0.5-fold and 2-fold, respectively. p-values were adjusted for false discovery rate using the Benjamini-Hochberg method. 6PG: 6-phosphoglycerate, αKG: alpha-ketoglutarate, R5P: ribose 5-phosphate, Xu5P: xylulose 5-phosphate, S7P : Sedoheptulose 7-phosphate, FBP: Fructose-1,6-bisphosphate.

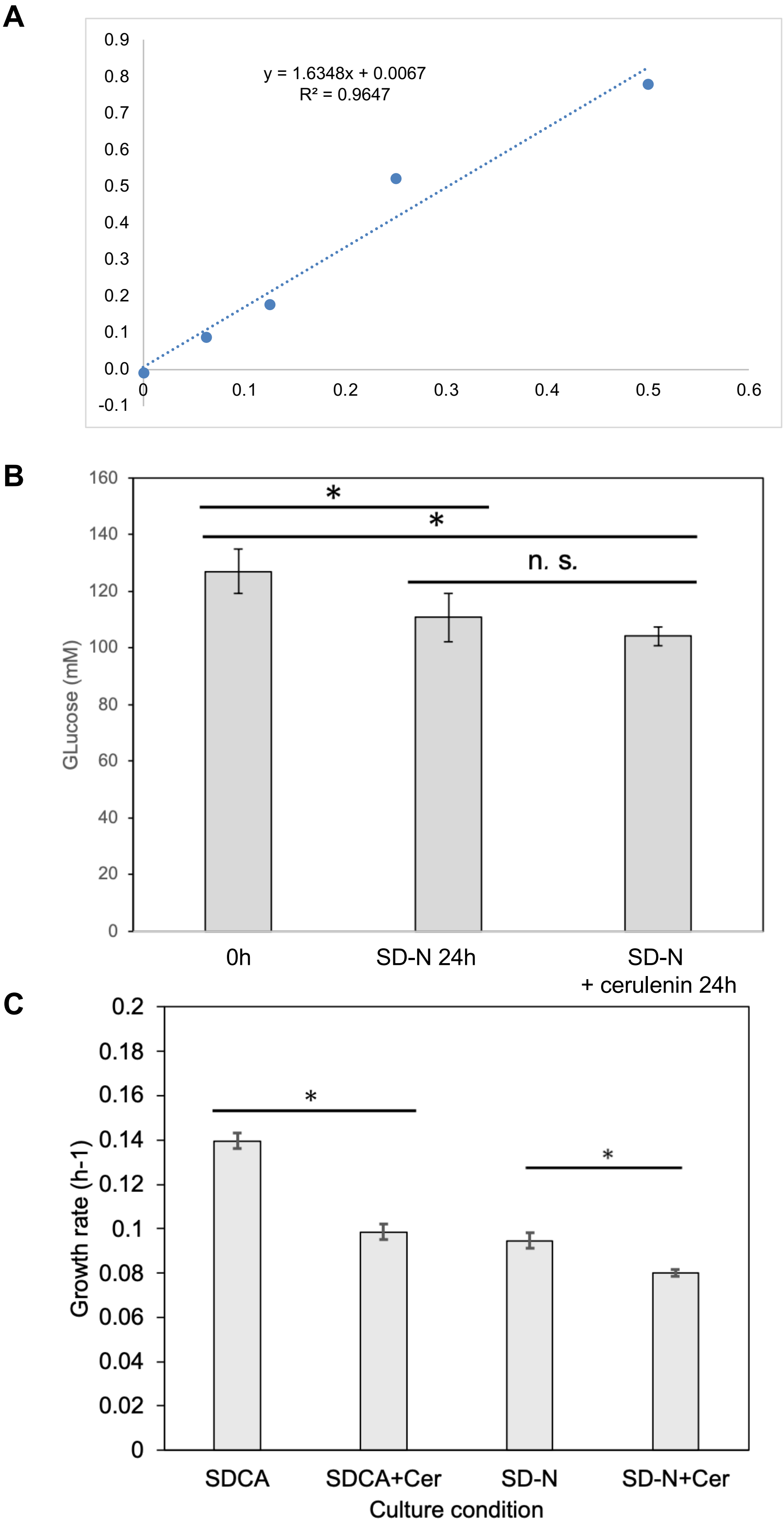
A
y = 1.6348x + 0.0067
R² = 0.9647
0.9
0.8
0.7
0.6
0.5
0.4
0.3
0.2
0.1
0.0
-0.1
0
0.1
0.2
0.3
0.4
0.5
0.6
B
GLucose (mM)
160
140
120
100
80
60
40
20
0
*
*
n. s.
0h
SD-N 24h
SD-N
+ cerulenin 24h
C
Growth rate (h-1)
0.2
0.18
0.16
0.14
0.12
0.1
0.08
0.06
0.04
0.02
0
*
*
SDCA
SDCA+Cer
SD-N
SD-N+Cer
Culture condition

A

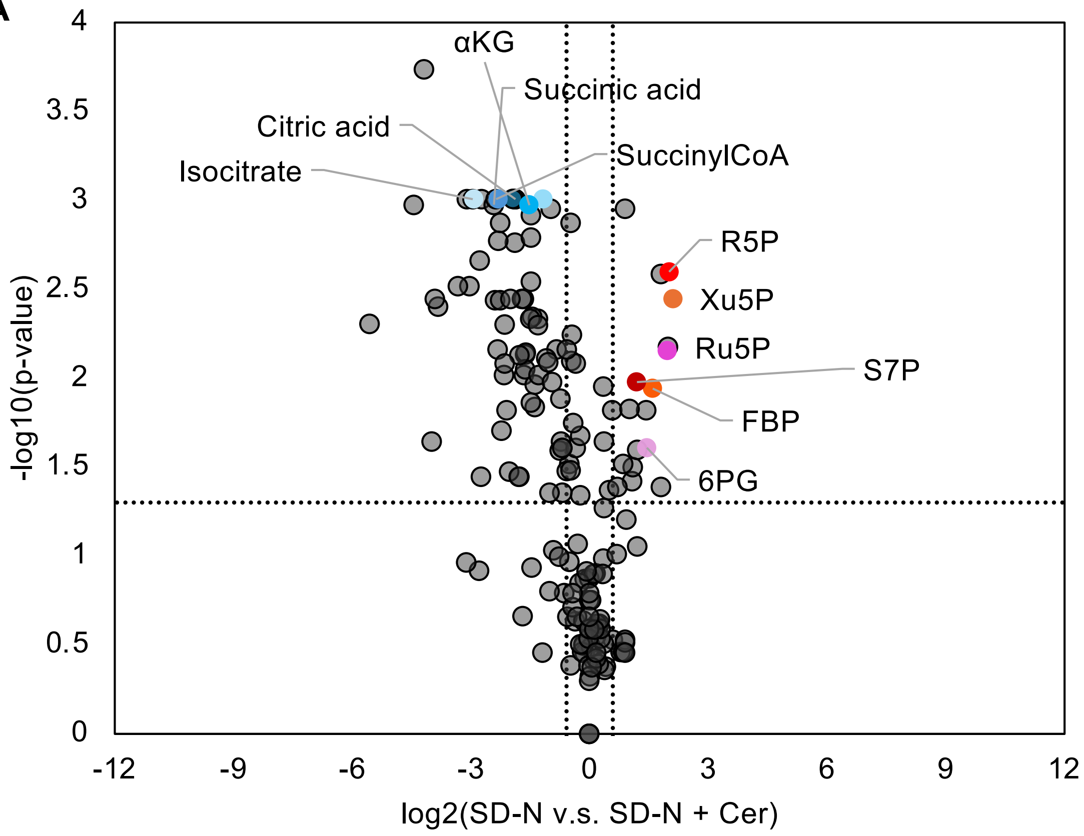
αKG
Succinic acid
Citric acid
SuccinylCoA
Isocitrate
R5P
Xu5P
Ru5P
S7P
FBP
6PG
-log10(p-value)
log2(SD-N v.s. SD-N + Cer)
4
3.5
3
2.5
2
1.5
1
0.5
0
-12
-9
-6
-3
0
3
6
9
12

B

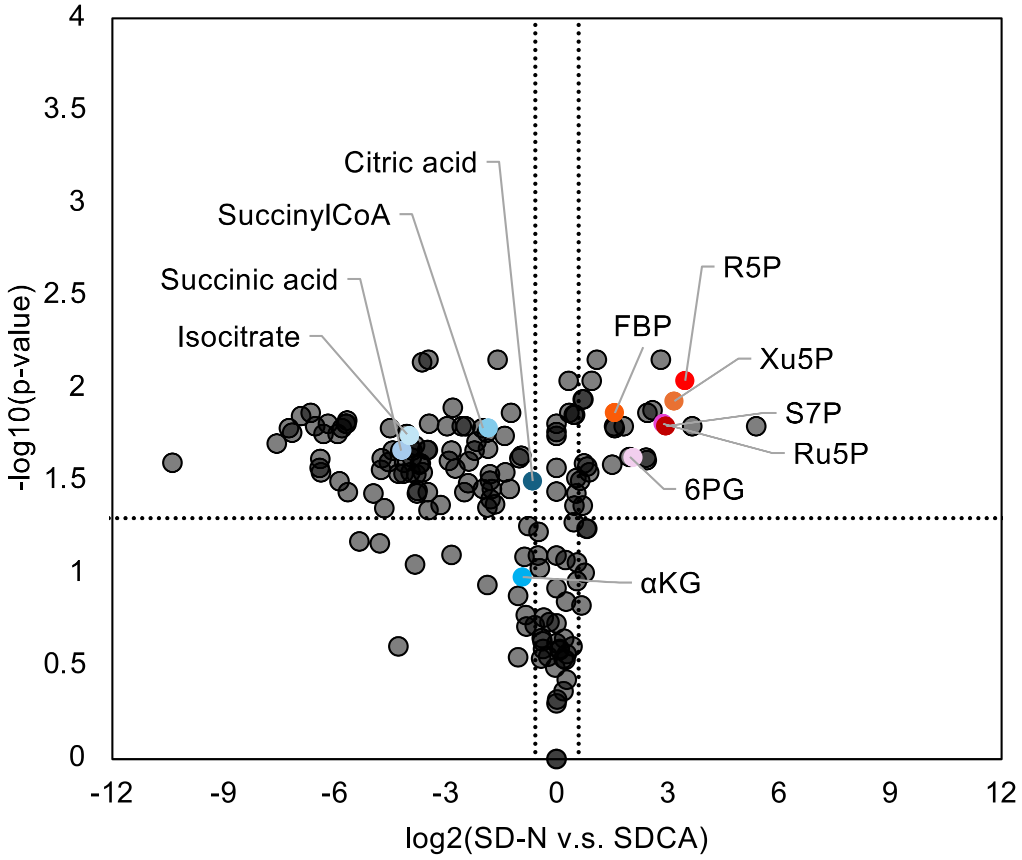
Citric acid
SuccinylCoA
Succinic acid
Isocitrate
R5P
FBP
Xu5P
S7P
Ru5P
6PG
αKG
-log10(p-value)
log2(SD-N v.s. SDCA)
4
3.5
3
2.5
2
1.5
1
0.5
0
-12
-9
-6
-3
0
3
6
9
12
